# Interest of circulating tumor DNA as a biomarker for canine cancers: illustration in histiocytic sarcoma, oral malignant melanoma and multicentric lymphoma

**DOI:** 10.1101/2020.07.10.189118

**Authors:** Anaïs Prouteau, Jérôme Alexandre Denis, Pauline De Fornel, Edouard Cadieu, Thomas Derrien, Camille Kergal, Nadine Botherel, Ronan Ulvé, Mélanie Rault, Amira Bouzidi, Romain François, Laetitia Dorso, Alexandra Lespagnol, Patrick Devauchelle, Jérôme Abadie, Catherine André, Benoît Hédan

## Abstract

Circulating tumor DNA (ctDNA) has become an attractive biomarker in human oncology and may be informative in cancer-affected dogs. By performing ddPCR or PARR methods, we detected tumor-specific point mutations, copy number alterations and chromosomal rearrangements in the plasma of cancer-affected dogs. It allowed the detection of ctDNA in 2/8 (25%) oral malignant melanoma cases, 12/13 (92.3%) lymphoma cases and 21/23 (91.3%) histiocytic sarcoma (HS) cases. The value of ctDNA to diagnose HS was explored in 133 dogs including 49 with HS. In this cohort, screening recurrent *PTPN11* mutations in plasma had a specificity of 98.8%, and a sensitivity between 42.8-77% according to HS clinical presentation, being higher in internal forms, especially with pulmonary location. Regarding lymphoma, the follow-up of four dogs showed that the minimal residual disease detection by targeting lymphoma-specific antigen receptor rearrangement in the plasma was concordant with the clinical evaluation. Moreover, ctDNA analysis appeared interesting to assess treatment response and to predict relapse.

This study shows that ctDNA is detectable in the plasma of cancer-affected dogs and is a relevant biomarker for diagnosis and clinical follow-up. With a growing interest in integrating natural canine tumors to explore new therapies, this biomarker appears promising in comparative oncology research.

## Introduction

Cell-free DNA (cfDNA) circulates in plasma and other body fluids such as urine, cerebrospinal fluid, pleural fluid and saliva as short double-stranded DNA fragments of about 160 base pairs. In healthy patients, cfDNA is released during cell deaths like apoptosis or necrosis, or by active secretion, and comes mainly from the hematopoietic lineage with minimal contributions from other tissues ^1–4^. Since its first discovery in human in 1948^5^, it was shown that cfDNA in the blood reflects various pathological processes such as inflammatory disease, sepsis, trauma, stroke and cancer, and its concentration is correlated with disease severity and prognosis ^2,6–8^. In cancer patients, a part of cfDNA is derived from tumor cells, and is referred to circulating tumor DNA (ctDNA). ctDNA carries tumor-related genetic and epigenetic alterations that are relevant to cancer development, progression and resistance to therapy ^8^.

In human medicine, next-generation sequencing (NGS) improved the knowledge of these genetic alterations and thus their identification in ctDNA. These alterations include for example mutations of tumor suppressor genes such as *TP53* in head and neck cancers ^9^, mutations in oncogenes such as *EGFR* or *KRAS* in non small cell lung cancer ^10,11^, copy number alterations (CNA), and chromosomal rearrangement like the one affecting the variable-diversity-joining (VDJ) receptor gene sequences in lymphoma

^12^. In the last decades, different strategies, including the study of loss of heterozygosity (LOH), DNA methylation, DNA integrity, NGS, or digital PCR were successfully developed to detect somatic alterations in patient’s plasma, making ctDNA a new powerful cancer biomarker ^8,13^. In addition to being a minimally invasive and robust approach, analysis of ctDNA has many clinical applications such as early cancer detection, prognosis, real-time monitoring of treatment response and identification of appropriate therapeutic targets and resistance mechanisms ^2,8,10,11,14^.

In veterinary medicine, cfDNA has recently gained attention and more particularly in dogs. As in human medicine, it has been found that cfDNA has a short half-life in canine plasma (around 5 hours) ^15^ and its concentration is increased in various diseases such as immune-mediated hemolytic anemia ^16^, sepsis, severe trauma or inflammation ^17–19^. In dogs, cfDNA concentration is also correlated with the severity of the disease and prognosis ^18^. It is also of high interest in canine cancers. It has been shown that dogs with lymphoid neoplasia and mammary carcinoma had a higher amount of cfDNA in plasma than controls ^19,20^, although this increase is not specific to cancer. Over the last 10 years, recurrent somatic alterations have been described in several canine cancers such as multicentric lymphoma ^21,22^, histiocytic sarcoma (HS) ^23–27^ and oral malignant melanoma (OMM) ^28–32^. Detection of such cancer-specific recurrent somatic alterations in ctDNA may allow to develop new minimally invasive biomarkers for diagnosis, prognosis and to assess treatment-response in veterinary medicine. In addition to those clinical considerations, tests based on ctDNA detection will be useful in the comparative oncology research field. Indeed, naturally occurring canine cancers have become relevant models for the study of rare human cancers, from the discovery of mutations to the development and the screening of targeted therapies ^27^.

The first objective of this study is to look for the presence of ctDNA and to confirm that several types of recurrent somatic alterations are detectable in the plasma of dogs with the example of three malignancies representing exceptional models for human counterpart: histiocytic sarcoma ^33^, multicentric lymphoma ^34^ and oral malignant melanoma ^35^. The second aim is to evaluate whether ctDNA can be used in veterinary medicine for diagnosis and response to treatment evaluation, using for proof of concept the diagnosis of histiocytic sarcoma and the monitoring of minimal residual disease in lymphoma-affected dogs treated by chemotherapy.

## Material and methods

### Sample collection

Blood and tissue biopsy samples from dogs were collected by veterinarians through the Cani-DNA BRC (http://dog-genetics.genouest.org) which is part of the CRB-Anim infrastructure (PIA1 2012-2022).

For each dog, between 2 to 5 ml of blood was taken during a routine examination or during the standard care pathway of the dog and was collected in ethylenediamine tetraacetic acid (EDTA) or Streck tubes (Streck cell-free DNA BCT genomax). The plasma was separated from blood cells and cell debris by centrifugations at 1500 g for 10 minutes and then 16 000 g for 10 minutes, and was stored at −20°C before DNA purification. This study working with canine samples was approved by the CNRS ethical board, France (35-238-13).

All the diagnoses of cancer were confirmed by histopathology or cytology by a board-certified pathologist (JA, LD).

### Case selection for monitoring of minimal residual disease (MRD)

The inclusion of dogs with a clinical follow-up to study minimal residual disease was done from October 2017 to October 2018. Dogs were prospectively enrolled in the study if they had a diagnosis of high-grade B-cell multicentric lymphoma confirmed by histopathology and if they started a chemotherapy protocol. Clinical staging was done on the basis of clinical examination, tomodensitometry and fine needle aspiration for detecting liver, spleen and thorax involvement, and myelogram for evaluating bone marrow infiltration. Evaluation of response to treatment and relapse was performed at each visit according to the criteria described for multicentric lymphoma by VCOG (Veterinary cooperative oncology group) ^36^, based on physical examination (lymph nodes measurement and appreciation of the clinician) and additional tests at the discretion of the clinician. Plasma was sampled before chemotherapy administration every week during the first month and then every 2-4 weeks until relapse or death.

### DNA extraction

Tumor DNA was extracted with DNA Nucleospin Tissue or FFPE Tissue DNA kits (Macherey Nagel, Düren, Germany), according to the manufacturer protocol, when RNAlater or FFPE tumor samples were available. Cell-free circulating DNA was extracted from 2 ml of plasma with NucleoSnap® DNA Plasma (Macherey Nagel, Düren, Germany), and eluted in 50µL of elution buffer according to the manufacturer’s protocol. When less than 2 ml of plasma was available, the volume was completed with PBS (Life Gibco).

### PCR for Antigen Receptor Rearrangements (PARR analysis)

For dogs with a diagnosis of high-grade multicentric lymphoma, paired samples (tumor DNA and plasma DNA) were analyzed by PCR for Antigen Receptor Rearrangement (PARR). M13-tailed primers were used to target immunoglobulin heavy chain genes for B lymphocytes and the T-cell receptor γ for T lymphocytes (Table1) ^18,37,38^. Amplification was achieved with the Type-it Multiplex PCR Master Mix (Qiagen), as previously described ^39^, and with the following conditions: denaturation at 92°C for 5 min, 35 cycles of 95 °C for 8 sec, 49 °C for 10 sec, and 72 °C for 15 sec (for IgH major, IgH minor and TCR Burnett), or with 30 cycles of 95°C for 45 sec, 48°C for 30 sec and 72 °C for 15 sec (for IgH Tamura, T Yagihara VA and Vb)^39^. The PCR amplicons were separated on a 370 ABI sequencer (Applied Biosystems) and Clonal profiles were determined with the Genemapper Software.

### Design of primers for digital droplet PCR (ddPCR)

ddPCR assays were designed using Biorad Droplet Digital PCR Assays and tools available online (https://www.bio-rad.com/digital-assays/#/assays-create/mutation) or copy number determination Biorad Droplet Digital PCR Assays (https://www.bio-rad.com/digital-assays/#/assays-create/cnd), for the detection of *PTPN11* mutations (E76K and G503V) or *MDM2, TRPM7*, CFA 9 copy number respectively. The region of CFA 9 was used as internal control as it was previously shown to have the higher stability of copy number in OMM.^28,40^ The Biorad **MIQE Context** locations of primers and probes are shown in Table 1.

**Table 1:**
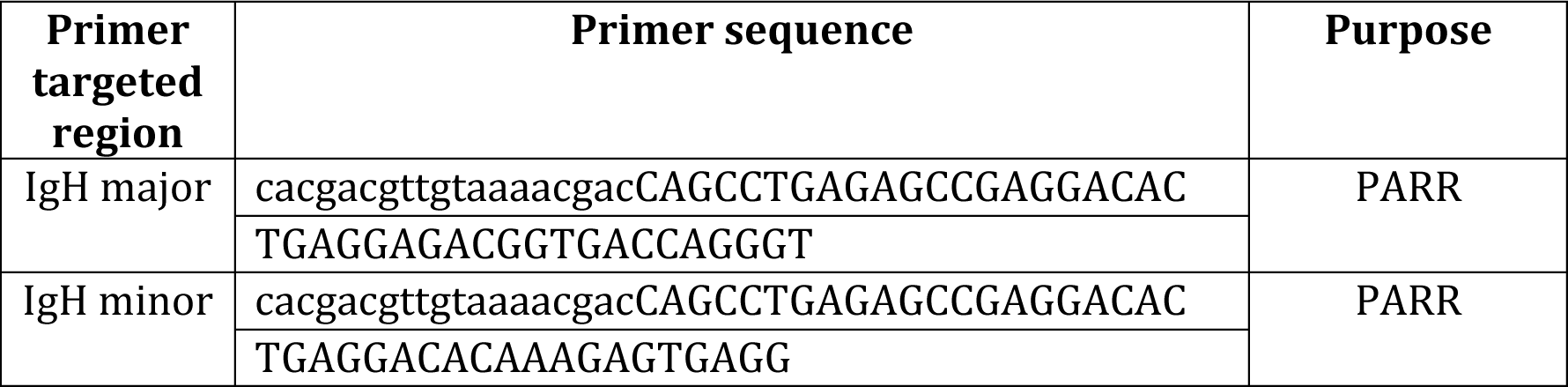
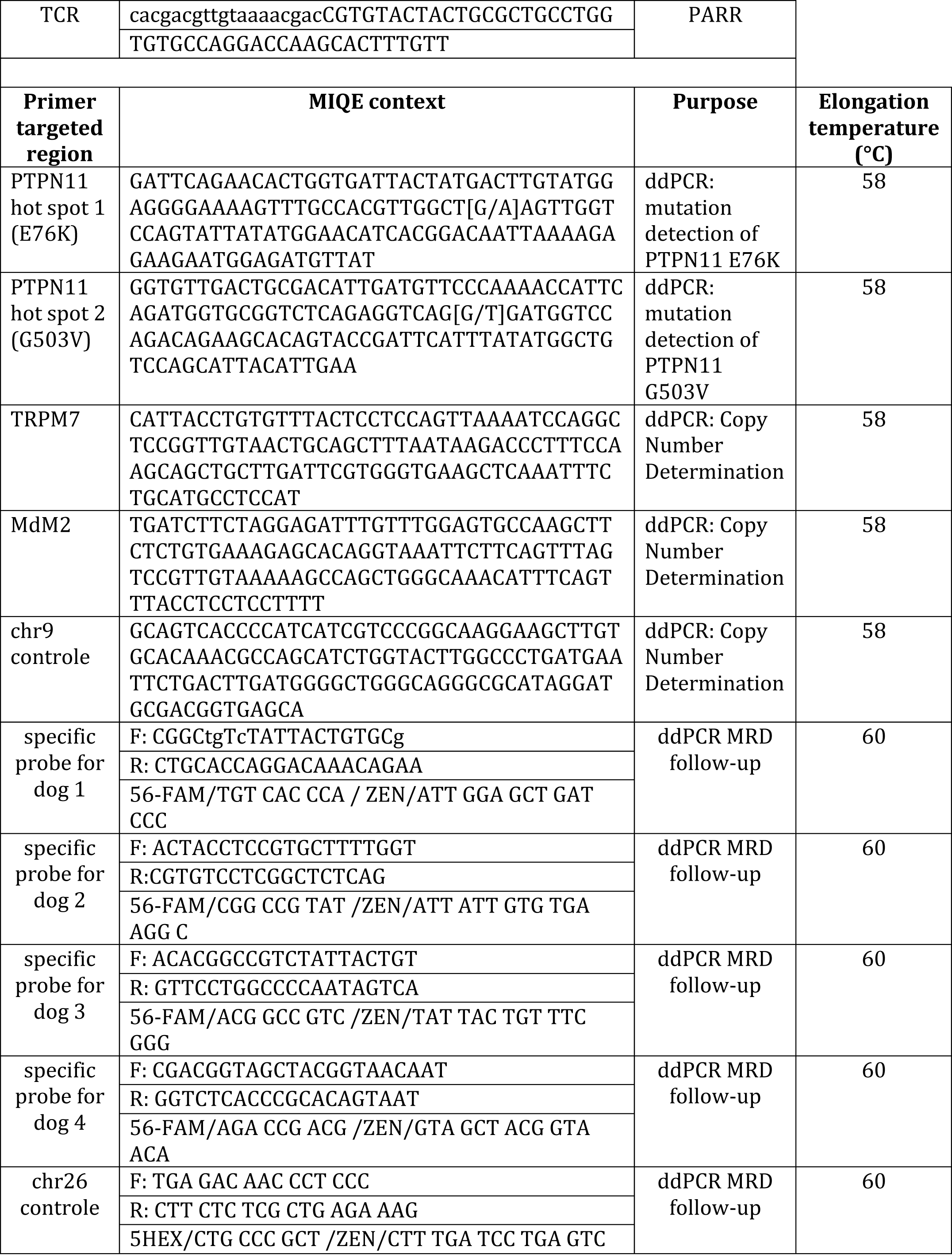
Primers used in the study.

For each dog with multicentric lymphoma examined for MRD, a PARR assay was performed on the tumor as described above, to isolate the clonal antigen receptor rearrangement. Then, the PCR product was sequenced with a 370 ABI sequencer using the BigDye Terminator v3.1 Cycle Sequencing Kit (Applied Biosystems). The sequences overlapping the CDR3 (Complementarity-determining region 3) were used to design clone specific primers and FAM fluorescent probes for ddPCR analysis, thanks to PrimerQuest Tool (IDT). A region on the CFA 26 was used as a control gene because it was previously found to be relatively stable in canine lymphomas ^41^ (Table1).

### ddPCR protocol and analysis

The ddPCRs assay comprised a total volume of 20 μL containing on the one hand the droplet Supermix (Bio-Rad Laboratories, Richmond, CA, USA) at final concentration of 500 nM of forward and reverse primers and 250 nM of VIC- and FAM-labeled probes and in the other hand the DNA input containing 8 µL of eluted cfDNA (mean: 15.2 ng) or 80 ng of tumor tissue DNA isolated from each patient sample. Next, the PCR reaction mixtures were partitioned into an emulsion of ∼20,000 droplets using a QX200™ ddPCR droplet generation system. PCR was conducted using the GeneAmp PCR system 9700 themocycler (Applied biosystems Foster City, CA, USA) using the following program: 95°C for 10 min; 40 cycles of 94°C for 30 sec and elongation temperature for 60 sec; 98°C for 10 min. Post PCR, droplets were analyzed on QX200™Droplet Reader (Bio-Rad). The concentrations of targeted sequences were calculated on the Poisson distribution using Quantasoft™ software Version 1.7.4 (Bio-Rad). Wild-type (wt) control DNA and non-template control reactions were included in each experiment. To ensure experiment quality, wells with total droplet counts of less than 8,000 were considered invalid and experiment was performed a second time to obtain a sufficient number of droplets.

*PTPN11* mutations frequency in cfDNA was determined by calculating the fraction of the *PTPN11* mutated molecule concentration A (copies/μl) to the *PTN11*wt reference molecule concentration B (copies/μl) (fraction= A/(A+B)). False positive rate estimation was determined by performing three experiments for each assay using the wild-type only samples, where total amounts of detected mutated positive droplets determined thresholds above which positive droplets in patient samples were to be considered as true positive. According to input quantity, only reactions containing at least one, two, three and four positive droplets for]0-7ng],]7-15ng],]15-100ng] and >100 ng input quantity respectively were classified as positive in the ddPCR analysis.

Concerning *MDM2* and *TRPM7* in canine OMM, copy number was determined using QuantaSoft analysis software (version 1.7.4, Bio-Rad) that calculated the ratio of the target molecule concentration A (copies/μl) to the reference molecule concentration B (copies/μl) multiplied by the number of reference copies in the canine genome (copy number = (A/B)*2). Those genes were considered amplified when the copy number was ≥ 3 and ratio’s confidence interval was > 2. The same method was used to quantify MRD in the plasma of lymphoma-affected dogs (A: lymphoma-specific antigen receptor rearrangement and B: CFA 26 region, control, and ctDNA% = (A/B)*100).

### Statistical analysis

Statistical analyses were performed using R studio software Version 1.1.463 (Vienna, Austria) ^42,43^.

## Results

1. Characteristics of the cohort

### Dog characteristics

A total of 133 dogs was included in the study, comprising 19 healthy dogs (mean age at sampling: 7.4 years old), 14 dogs with non-cancerous diseases (mean age at sampling: 7.3 years old) and 100 cancer-bearing dogs (mean age at sampling: 8.2 years old) (Supplementary Table 1). Among healthy dogs and dogs with cancers, the most frequent breeds were Bernese mountain dog (9/19 and 45/100 respectively), and flat-coated retriever (n=8/19 and 7/100 respectively). The most represented cancer in our study was histiocytic sarcoma (n=49), with 17 disseminated forms, 30 localized forms and 2 unknown (Supplementary Table 1). There were 16 oral malignant melanomas, 25 multicentric lymphomas (21 high-grade and 3 low-grade lymphomas), and 10 other cancers including ocular melanoma (n=2), soft tissue sarcoma (n=2), nephroblastoma (n=1), hemangiosarcoma (n=2) hepatocellular carcinoma (n=1), Langerhans cell sarcoma (n=1) and glioma (n=1).

### Quantification of cell-free DNA (cfDNA) in plasma

cfDNA in the plasma of all dogs was accurately quantified in the 133 dogs by the ddPCR method using the *PTPN11*wt gene as a reference. The median cfDNA concentration in healthy, HS, multicentric lymphoma, OMM, other cancers and non-cancerous diseased dogs were 339ng/ml (mean 678ng/ml; range 6.7-3895), 240ng/ml (mean 827ng/ml; range 8.1-6320), 1168ng/ml (mean 4764ng/ml; range 25.4-35079), 106ng/ml (mean 3833ng/ml; range 8.2-49910), 185ng/ml (mean 1282ng/ml; range 19.6-10746) and 259ng/ml (mean 454ng/ml; range 0.7-3030) respectively (Figure 1). Dogs with multicentric lymphoma had a significantly higher cfDNA concentration in plasma than healthy dogs (Wilcoxon test, p=0.034), dogs with HS (Wilcoxon test, p=0.005), dogs with OMM (Wilcoxon test, p=0.009), dogs with other cancers (Wilcoxon test, p=0.025) and dogs with non-cancerous diseases (Wilcoxon test, p=0.006). There was no significant difference in cfDNA concentration between the other groups of dogs, particularly between healthy dogs and dogs with other cancers than multicentric lymphoma. Thus, this illustrates that cfDNA is not a sensitive marker for tumor diagnosis and that the detection of tumor specific genetic alterations is required to confirm the existence of a malignancy.

**Figure 1:**
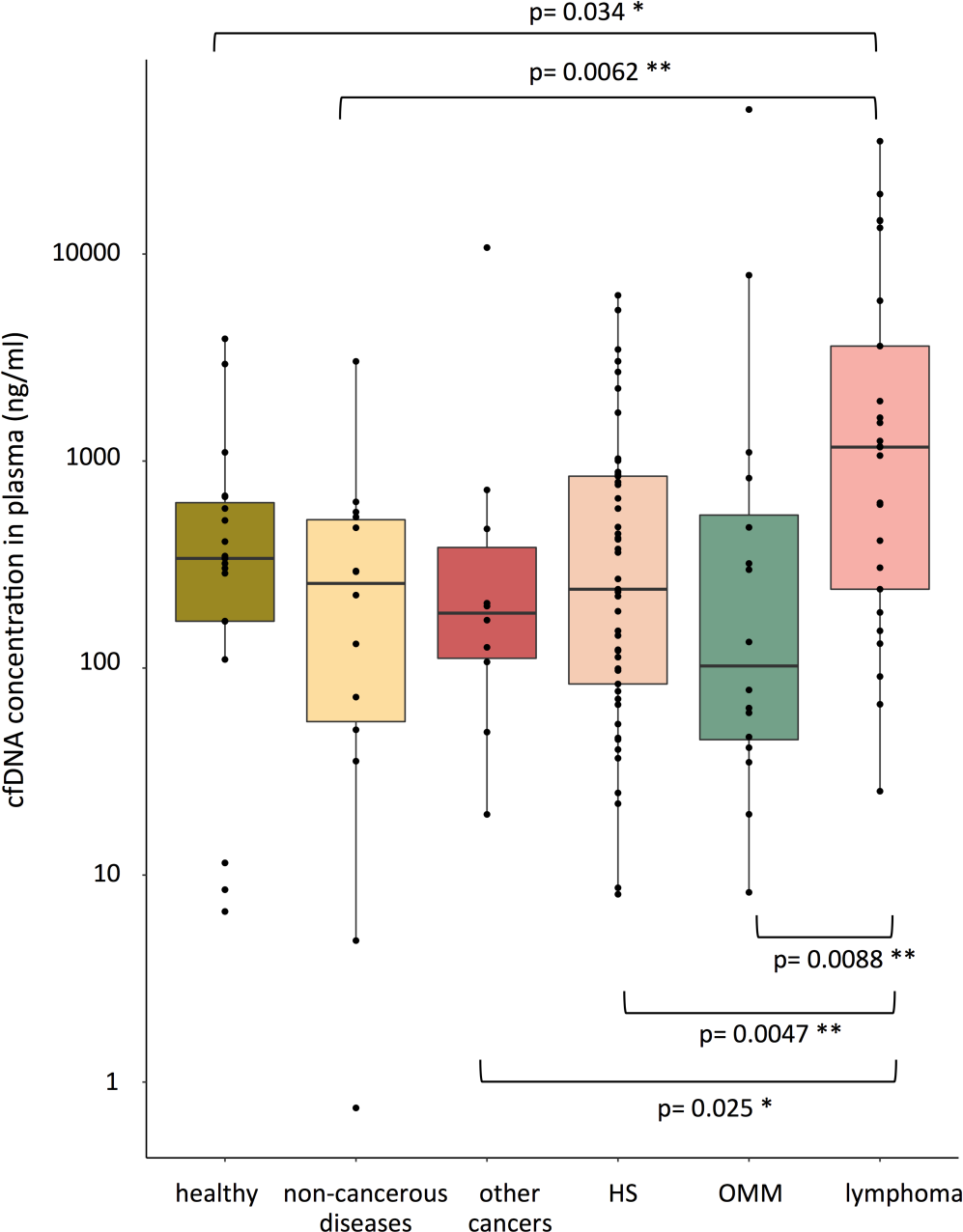
Quantification of cfDNA (ng/ml) in the plasma of dogs under different conditions: healthy, non-cancerous disease, various cancers, histiocytic sarcoma (HS), oral malignant melanoma (OMM) and multicentric lymphoma. The amount of DNA was determined by Poisson‘s law.

2. Detection of ctDNA in plasma of dogs bearing cancer

### Detection of tumor specific mutation in the plasma of dogs with HS

The knowledge of tumor-specific and recurrent hotspot of mutation is required to use a point mutation in the screening of ctDNA. *PTPN11* is an oncogene involved in the MAPKinase pathway and is mutated in 23 to 56.7% of HS cases, mainly in two hotspot (E76K or G503V), depending on clinical presentation ^25–27^.

In this study, *PTPN11* mutation was found in 23/45 HS tumor samples. The same mutation was detected in the plasma of 21/23 (91.3%) corresponding dogs, and the fraction of ctDNA in overall cfDNA varied from 0.056 to 36% (Supplementary Table 1). Moreover the mutations were not detected in any plasma from dogs with *PTPN11*wt tumors (Figure 2).

**Figure 2:**
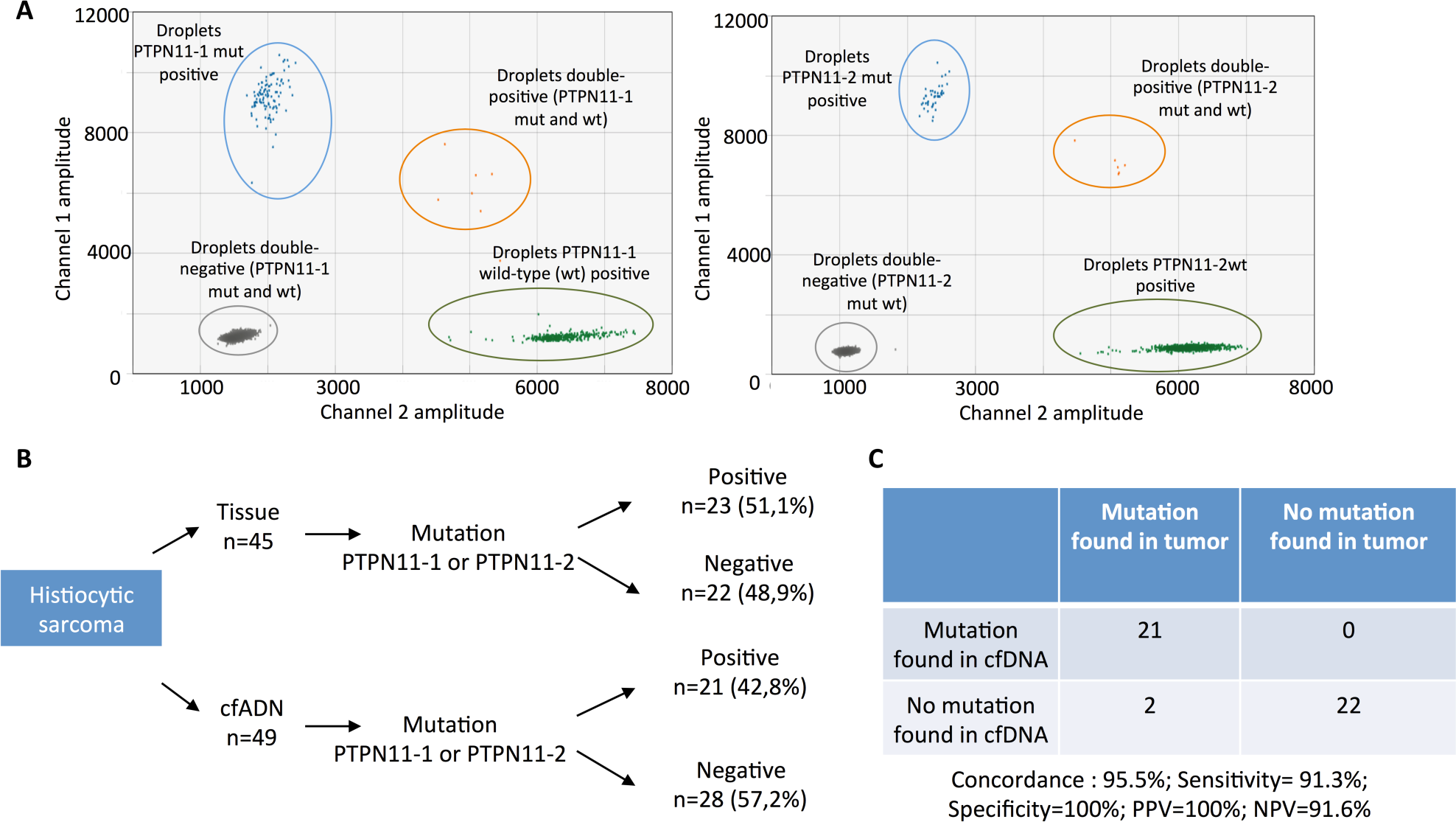
**A**: Example of data output from our ddPCR multiplex assay for absolute quantification of *PTPN11* mutations (PTPN11-1 for E76K and PTPN11-2 for G503V) in canine HS. Exemplary two-dimensional cluster plots in which PTPN11-1mut (left) or PTPN11-2mut (right) are plotted against PTPN11-1 wild-type (wt) and PTPN11-2wt respectively. Droplets form clusters that should arrange orthogonally to each other and represent PTPmut/PTPwt negative (double-negative droplets, grey), PTPmut positive (blue), PTPwt positive (green), and PTPmut/PTPwt positive (double-positive droplets, orange). **B**: Flowchart summarizing results of the ddPCR assay targeting *PTPN11* mutations (E76K and G503V)on tumor and plasma samples of 49 dogs with HS. cfDNA: cell-free DNA. **C**: Parameters of the test for *PTPN11* mutation detection in plasma of dogs with HS. PPV: positive predictive value, NPV: negative predictive value.

### Detection of chromosomal rearrangement in the plasma of dogs with lymphoma

ctDNA can be detected also by the identification of tumor-specific chromosomal rearrangements, as performed in human affected by multicentric lymphomas.^12^ VDJ recombination of immunoglobulin and T-cell receptor occurs in early stage of lymphocyte maturation and is specific to the lymphocyte clone. Thus, in lymphomas, the VDJ sequence is unique and constitutes a marker of the tumor clone ^12^. In order to detect lymphoma-specific VDJ rearrangement, we performed PCR for antigen receptor rearrangement (PARR) in 14 dogs with high-grade multicentric lymphoma. Immunophenotype was available for 10/14 dogs and all of them were B-cell lymphomas. PARR was positive and detected a clonal rearrangement in the tumor of 13/14 dogs. To detect ctDNA, PARR was performed on cfDNA extracted from plasma at the time of diagnosis. All but one sample were positive and presented the same clone in tumor and plasma samples (Figure 3).

**Figure 3:**
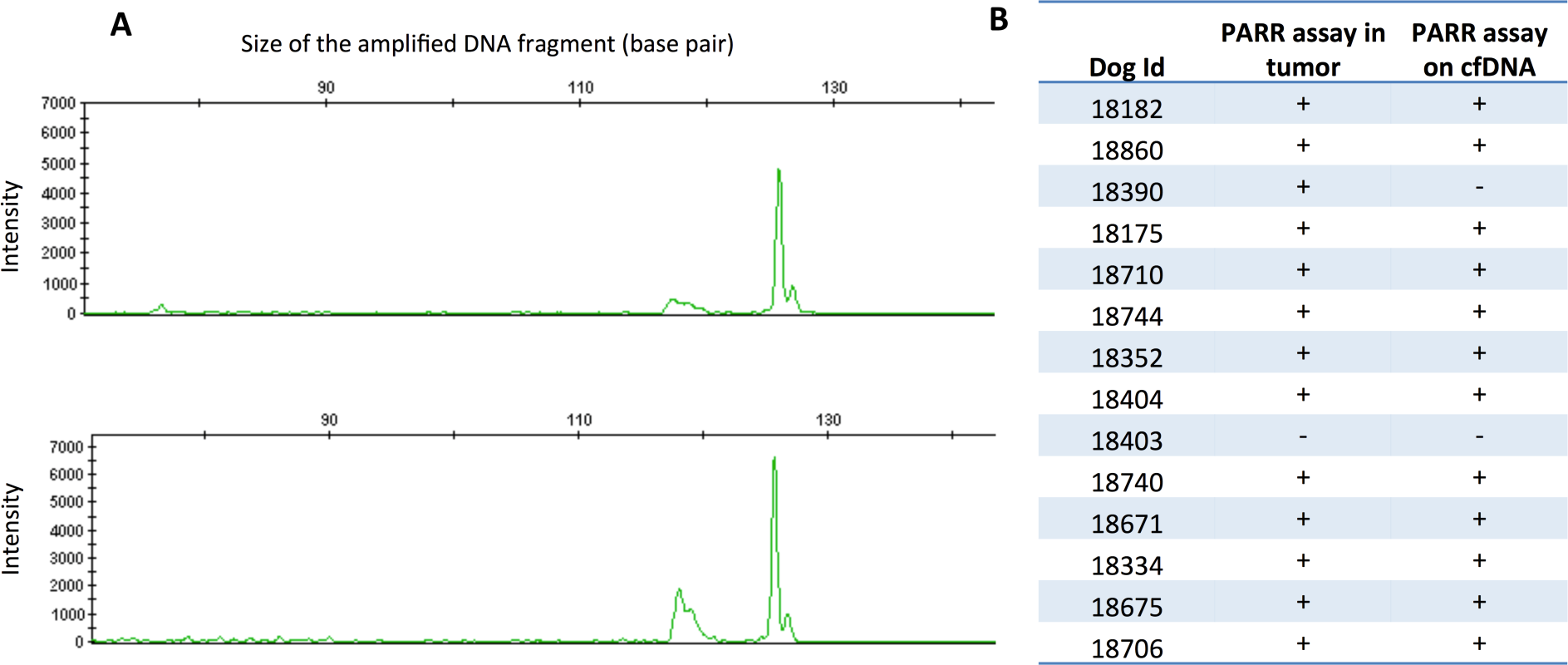
**A**: Results of a PARR genotyping on DNA extracted from a tumor sample (above) and from plasma (below) of a dog with high-grade multicentric lymphoma (Id 18175). The same clonal antigen receptor rearrangement (IgH major) was found with a high intensity in the tumor and in plasma. **B**: Results of our PARR assay on canine lymphoma. We report the results on DNA from tumor and DNA extracted from plasma. PARR was positive (clonal band) in tumor and cfDNA for 12/13 dogs.

These results illustrate that tumor-specific chromosomal rearrangement are detectable in the plasma of dogs with multicentric lymphomas and that a majority of canine lymphoma patients (92.3% in our cohort) present ctDNA at the time of diagnosis.

### Detection of copy number alterations (CNAs) in the plasma of dogs with OMM

As a proof of concept that CNAs are detectable in the plasma of dogs bearing cancer, we focused on OMM. In order to detect ctDNA in OMM patients, we targeted *MDM2* and *TRPM7* amplifications –present in around 50% of canine OMM-^28,30–32^. Copy number was determined in tumor and plasma by ddPCR. On the 10 OMM cases with available tumor samples, amplification of *MDM2* was found in 3/10 tumors, and was detected in the plasma of 1/3 corresponding dog (Table 2). On the other hand, amplification of *TRPM7* was found in 8/10 tumors, and was detected in the plasma in only one case (Table 2).

**Table 2:** Results of the ddPCR assay to detect copy number alterations (CNA) of *TRPM7* and *MDM2* on tumor and plasma of dogs with oral melanoma. Eight dogs had at least one gene amplified in the tumor, and in two cases, a copy number imbalance was found in the plasma.

|  | Tumor | cfDNA |
| --- | --- | --- |
| Dog Id | Gain detected: copy number (95% confidence interval) | Gain detected: copy number (95% confidence interval) |
| 19006 | *TRPM7*: 37,90 (28,41-47,39) | *TRPM7*: 2,80 (2,58-3,03) |
| 18394 | *TRPM7*: 7,98 (7,71-8,25) | No |
| 18657 | *TRPM7*: 3,01 (2,89-3,14) | No |
| 18718 | *MDM2*: 21,26 (19,37-23,15) | *MDM2* : 3,02 (2,19-3,85) |
|  | *TRPM7*: 7,46 (7,17-7,75) | No |
| 18395 | *MDM2*: 52,96 (45,74-60,17) | No |
|  | *TRPM7*: 8,78 (8,37-9,19) | No |
| 18303 | *TRPM7*: 8,88 (8,55-9,20) | No |
| 18333 | *MDM2*: 4,30 (3,91-4,68) | No |
|  | *TRPM7*: 27,24 (25,94-28,54) | No |
| 18848 | *TRPM7*: 9,12 (8,78-9,46) | No |

Altogether, those results show that specific tumor CNAs were detectable in the plasma of 2/8 (25%) dogs affected by OMM, proving that part of circulating DNA is derived from tumor cells in at least a proportion of this cancer.

3.Illustration of potential clinical application for veterinary medicine and oncology research

ctDNA has become an attractive biomarker to monitor human cancer patients, with many clinical applications such as early cancer detection, prognosis, real-time monitoring of treatment response and identification of appropriate therapeutic targets and resistance mechanisms. In the same time, dog is recognized as underutilized model to develop new therapies for high-grade B-cell lymphoma ^34,44^ or rare cancers such as HS ^33^. Here we illustrate two potential applications for veterinary medicine and comparative oncology: the diagnosis and the monitoring of minimal residual disease.

### Interest of ctDNA for diagnosis: example of *<u>PTPN11</u>*<u>mutation in canine HS</u>

The knowledge of a recurrent tumor-specific mutation can be also useful to develop a non-invasive molecular diagnostic tool ^45,46^.

In order to assess the sensitivity and specificity of *PTPN11* mutation screening in plasma to diagnose HS, we tested the plasma of 49 dogs with HS, 19 healthy dogs, 14 dogs with non-cancerous diseases and 51 dogs with various cancers (multicentric lymphoma, OMM, soft tissue sarcoma, Langerhans cell sarcoma, nephroblastoma, hemangiosarcoma and ocular melanoma) (Supplementary Table 1). The *PTPN11* mutation was detected in the plasma of 21/49 dogs with HS, and 1 dog with a non-cancerous disease. It was not found in the plasma of healthy dogs and dogs suffering from other cancers. Considering those results, the screening of *PTPN11* mutation in the plasma had a specificity of 98.8% and a sensitivity of 42.8% for the diagnosis of canine HS. However, the sensitivity varied according to HS characteristics and clinical presentations, and was higher among Bernese mountain dogs (53.8%), in cases of disseminated forms (58.8%), internal organs involvement (59.4%) and especially with thoracic location (77%) (Supplementary Table 2). This is due to the higher frequency of *PTPN11* mutations in Bernese mountain dogs (Fisher exact test, p=0.0015), internal clinical presentations (Fisher exact test, p=0.008) and when there is a pulmonary involvement (Fisher exact test, p=0.002) in this cohort.

### Interest of ctDNA for monitoring minimal residual disease (MRD): example of high-grade B-cell multicentric lymphoma

In lymphoma, the unique VDJ rearrangement of the tumor clone constitutes a “barcode” that is used for the follow-up of human lymphoma patients in clinical trials ^12^.

Four dogs presenting a high-grade B-cell multicentric lymphoma were prospectively included and their lymphoma-specific VDJ rearrangements were sequenced to further perform ctDNA analysis during follow-up. Three dogs received a cyclophosphamide, doxorubicin, vincristine, plus prednisolone (CHOP)-^47–49^ based chemotherapy and one dog (dog 3) received only vincristine injections. L-asparaginase was used in two cases at induction. In the case of absence of response or relapse, L-asparaginase and lomustine were used as rescue treatments. Before initiation of chemotherapy, ctDNA was detected in all cases and was determined as a ratio (%) of cfDNA (mean proportion: 15.5% of total cfDNA before initiation of chemotherapy). The monitoring details of each case are summarized below and represented in Figure 4.

**Figure 4:**
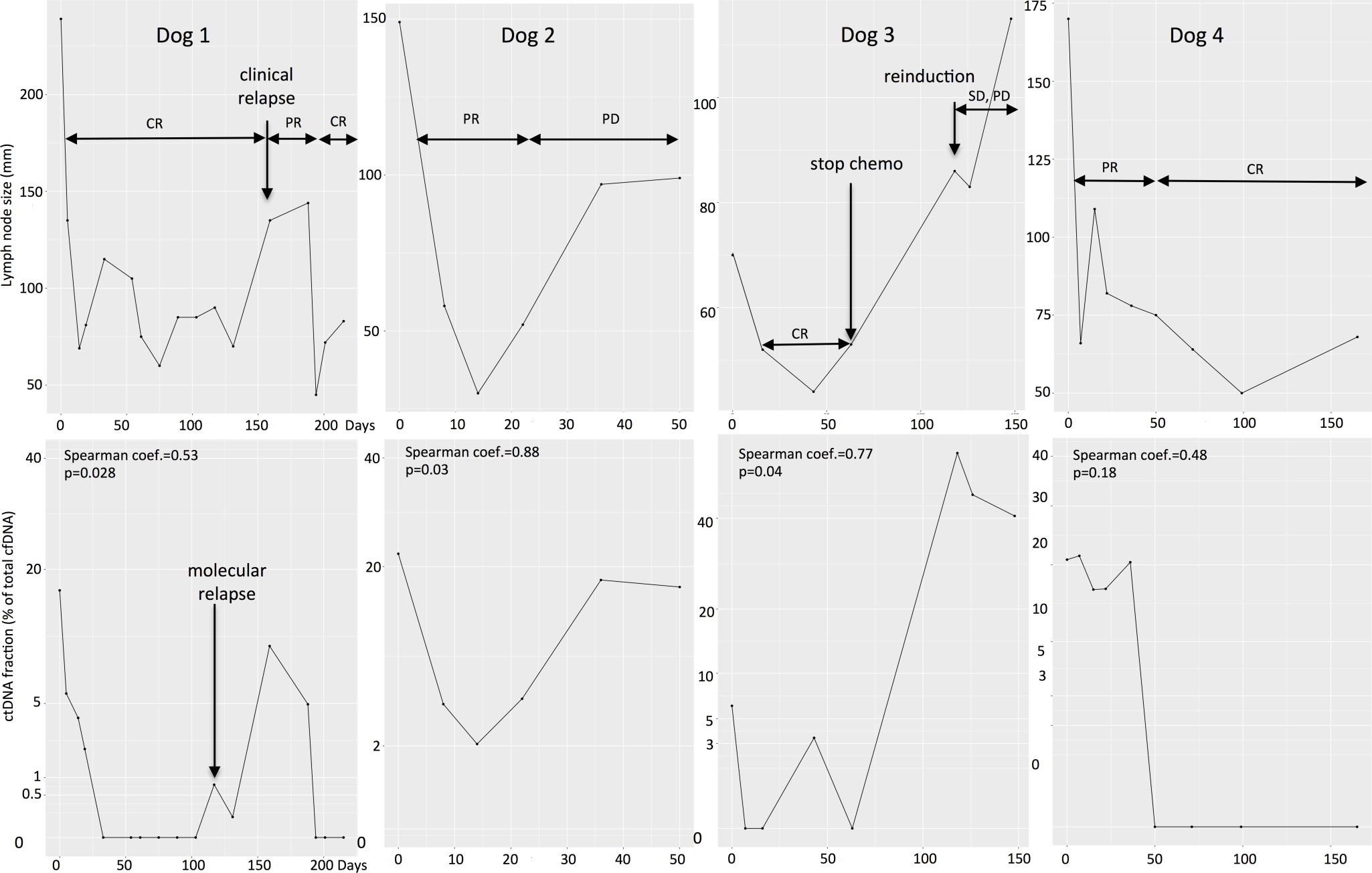
Course of the disease of four lymphoma-affected dogs treated by chemotherapy (dog 1: 18182, dog 2: 18710, dog 3: 18680, dog 4: 18744). For each dog, the figures above represent tumor burden with lymph node size (sum of longest diameter of at least 2 lymph nodes, in mm) and appreciation of the clinician. The figures below represent ctDNA measurement (% of total cfDNA in plasma) during disease course. For each case, a Spearman correlation was done to assess correlation between tumor burden and ctDNA quantity (coefficient and p-value). CR: complete response, PR: partial response, SD: stable disease, PD: progressive disease

Dog 1: This dog had a stage IV multicentric lymphoma and presented 17% of ctDNA in plasma before initiation of chemotherapy. A complete response (CR) was achieved 14 days after initiation of chemotherapy while molecular remission measured by ddPCR (ie. no ctDNA detected) occurred at day 33. At day 117 ctDNA was detected again in the plasma in a very small proportion (0.77%). The clinical relapse was objectified at day 159, ie 42 days after ctDNA detection. After reinduction, a second complete response was attained with a matched molecular remission. At day 274, the owner decided to stop chemotherapy. During disease course, ctDNA quantity and lymph node size were significantly positively correlated (Spearman correlation test, coefficient= 0.53, p=0.028).

Dog 2: This dog had a stage V multicentric lymphoma and we quantified 22% of ctDNA in its plasma before initiation of chemotherapy. The dog had only a partial response (PR) to therapy and was euthanized at day 50 after initiation because of disease progression (PD). Molecular monitoring fitted clinical observation as ctDNA was detected all the time during disease course. Lymph node size and ctDNA quantity were positively and significantly correlated (Spearman correlation test, coefficient= 0.88, p=0.03).

Dog 3: This dog had a stage IV multicentric lymphoma and presented only 6.25% of ctDNA in plasma before chemotherapy initiation. This lower ctDNA fraction may be due to a prednisolone treatment that was started 7 days before the beginning of the CHOP chemotherapy protocol. The dog experienced a CR at day 7 that matched with molecular remission. ctDNA was detected again at day 43 whereas the dogs was still in CR, but was not detected at the next sampling (day 63). The owner decided to stop chemotherapy at day 63 while the dog was in CR and molecular remission. One month after, the tumor recurred and a second chemotherapy was initiated but without significant response (stable disease –SD-followed by PD). This clinical evolution matched with the detection of high amount of ctDNA. The dog died one month after initiation of the second chemotherapy. Tumor burden and ctDNA quantity were significantly and positively correlated (Spearman correlation test, coefficient= 0.77, p=0.04).

Dog 4: This dog attained a CR at day 50 after chemotherapy initiation and the owner decided to stop the treatment at day 165 while the dog was still in CR. The complete clinical response was matched with a molecular remission that lasted until the dog was lost to follow-up. In that case the correlation between lymph node size and ctDNA quantity was positive but not significant (Spearman correlation test, coefficient= 0.48, p=0.18).

Altogether, those results suggest that the ctDNA kinetic during a chemotherapy treatment in dogs affected by multicentric lymphoma is correlated with lymph node size and is reflective of treatment response. Moreover, ctDNA could be more accurate than lymph node measurement as we were able to detect a molecular relapse 42 days before the clinical relapse.

## Discussion

cfDNA is an attractive biomarker in human medicine and particularly in oncology. Thus, it has also been evaluated in different canine diseases ^16,17,19,20,50–53^. As in human, cfDNA concentration in dogs is correlated with the severity of the disease and prognosis ^16,17,20,53^. Consistently with previous studies, we found that dogs with multicentric lymphoma had significantly higher concentration of total cfDNA than healthy dogs, dogs with non-cancerous diseases and dogs with other cancers ^20,51^. Our cfDNA concentrations were in agreement with those of previous studies, although results were highly variable in terms of median cfDNA concentrations, ranging from 4 to 583ng/ml in healthy dogs for example. In those studies, authors used different extraction kits and quantification methods, ^16,20,51,53^ that could in part explain this inter-study variability. Thus, this high variability of cfDNA concentration in healthy dogs limits its interest as a global biomarker for cancer diagnosis.

In human oncology, current studies focus on the analysis of DNA released from tumor cells in plasma (ctDNA), which is much more cancer-specific. Today one ctDNA assay for the detection of *EGFR* mutation in patients with non-small cell lung cancer (NSCLC) has been approved by the Food and Drug Administration, and ctDNA assays for *EGFR* in NSCLC and *KRAS* in colorectal cancer are available for commercial use in Europe ^3,4,54^. As in human, ctDNA has also been found in some canine cancers such as lymphoma ^20^, mammary carcinoma ^50^ and more recently in pulmonary adenocarcinoma ^52^. While ctDNA can be released in many biological fluids, as it was demonstrated with *BRAF* mutation detection in urine for the diagnosis of canine urothelial and prostatic carcinomas ^55^, we focused on the interest of plasma as liquid biopsy in the present study. We showed that ctDNA can be detected in canine histiocytic sarcoma, oral malignant melanoma and multicentric lymphoma. Further studies are needed to confirm those findings in larger cohorts, and to explore the potential association between ctDNA and clinical parameters. However, the high frequency of ctDNA detection among HS and lymphoma cases suggests that ctDNA is a promising biomarker in those hematopoietic malignancies.

We identified ctDNA thanks to several tumor-associated alterations and showed that point mutations, chromosomal rearrangements and CNAs were detectable in the plasma of dogs with cancer. According to the targeted alterations, we used different methods, with their advantages and drawbacks. It is important to highlight that the performance of ctDNA detection largely depends on the level of ctDNA in the patient’s plasma and the sensitivity of the method used. Since the release of a lot of cfDNA occurs in case of lymphoma (median cfDNA: 1168 ng/ml in the present study), it is not surprising that the PARR assay was able to detect ctDNA in 92.3% (12/13) of lymphoma affected dogs at the time of diagnosis, confirming a previous study that found ctDNA in 78% (7/9) of dogs with multicentric lymphoma^20^ using the same method. In dogs with HS, the cfDNA concentration was much lower (median 240 ng/ml), but as we used a highly sensitive method (ddPCR), we were able to detect *PTPN11* mutation in the plasma of 91.3% (21/23) of dogs with *PTPN11* mutated tumor. This assay had positive and negative predictive values of 100% and 91.6% respectively to detect *PTPN11* mutation. Recently, Lorch et al. used the same method to search *HER2* mutation in matched tumor and plasma samples of dogs with pulmonary adenocarcinoma ^52^. They found the mutation in only 33% of dogs with *HER2* mutated tumor. This low detection rate compared to our results in dogs with HS is probably due to the lower fraction of ctDNA released in pulmonary adenocarcinoma compared to HS, and thus the lower DNA amount available for mutation detection (median cfDNA of 3.7 ng/ml of plasma in dogs with pulmonary adenocarcinoma versus 240 ng/ml in dogs with HS). Low cfDNA amount was also an issue when we studied ctDNA in OMM. The most recurrent genetic alterations in that cancer are CNAs targeting oncogenes like *MDM2* and *TRPM7* ^28,30–32^, and we detected those tumor-associated amplifications in cfDNA with ddPCR in only 1/3 and 1/8 cases respectively. Detecting CNAs in cfDNA is challenging, and depends also on the level of amplification of the gene in the tumor which in turns, involves deeper sequencing ^56^. In humans, more reliable methods are used to detect CNA in cfDNA like low-pass whole genome sequencing ^57^. Beck et al. used whole genome sequencing on tumor and plasma samples of 2 dogs with mammary carcinoma and found a good concordance of tumor and cfDNA copy number profiles for the regions with high-level amplifications ^50^. However, this method requires high sequencing coverage and would not yet be cost-effective for veterinary medicine.

Another objective of this study was to evaluate the potential clinical application of ctDNA analysis in veterinary oncology. Two aspects were investigated: the value of ctDNA analysis as a tool for HS diagnosis, and for MRD follow-up in lymphoma-affected dogs. Diagnosis of canine histiocytic sarcoma is sometimes challenging because of hidden tumor locations (e.g. lungs, mediastinum, central nervous system…) and the potential difficulty to differentiate it from reactive inflammatory disease or other malignancies ^58^. We thus assessed the value of a ctDNA test in order to diagnose HS using a non-invasive method. In our study population of 133 dogs, screening *PTPN11* in plasma for HS diagnosis had a specificity, sensitivity, positive and negative predictive values of 98.8%, 42.8%, 95.5% and 74.8% respectively. We found that *PTPN11* mutation was highly specific of HS, as it was never detected in the plasma of dogs with other cancers. Only one dog was classified as false positive and concerned a 12 years-old Bernese mountain dog sampled at the time of euthanasia because of old-age, and who was classified in the “non-cancerous diseases” group. Two hypotheses can explain this false positive result. First, this old dog could suffer from an undiagnosed or early stage HS, and unfortunately, necropsy was not performed to confirm this hypothesis. Second, somatic variants associated to hematologic cancers have been found in the plasma of apparently healthy people, due to “clonal hematopoiesis” ^4^. Clonal hematopoiesis refers to the selective expansion of hematopoietic stem cells with somatic mutations of genes involved in hematologic malignancies. In human, it increases with age and it is correlated with the risk to further develop a hematologic cancer. This has not been investigated in dogs yet, and further studies are needed in predisposed breeds to look for *PTPN11* mutation in the plasma of apparently healthy dogs to see if clonal hematopoiesis precedes HS development. The relatively low sensitivity of this test is due to the fact that *PTPN11* is not mutated in all HS cases (i.e. 50% of cases in this study) and is found especially in Bernese mountain dogs ^25^ and in the disseminated forms ^27^. In the present study, all mutated cases were Bernese mountain dogs. Moreover, we targeted only the two most frequent *PTPN11* mutations (E76K and G503V), while some HS cases present other *PTPN11* mutations ^27,59^. Recently, mutations of *KRAS* and *BRAF* were also found in 3 to 7% and less than 1% of canine HS respectively, and were found to be exclusive of *PTPN11* mutation ^25,27^. Thus, the sensitivity of this diagnostic test could be improved by integrating these other mutations of *PTPN11, KRAS* and *BRAF* in the ddPCR assay. The best sensitivity was obtained in the subgroup of dogs with visceral form and pulmonary involvement (10/13, 77%). Thus, screening of *PTPN11* mutations in plasma would be interesting to confirm a clinical suspicion of HS especially in case of pulmonary involvement and in Bernese mountain dogs. Further studies are needed to define the value of this test in a population including more breeds and to explore the potential interest of this biomarker for early stage diagnosis and for treatment follow-up in dogs with HS. In the future, this test could be considered in veterinary oncology, to guide treatment procedure toward targeted therapy. Indeed, others and us showed that *PTPN11* mutations were associated with the activation of the MAPK pathway, and *in vitro* test showed that canine HS cell lines were sensitive to MEK inhibitors ^27,59,60^. As our ctDNA assay demonstrated a sensitivity of 91.3% and a specificity of 100% for the detection of *PTPN11* mutation in the plasma of dogs carrying *PTPN11* mutated HS, it could efficiently select dogs susceptible to benefit from this targeted therapy. Finally, since the same mutations of *PTPN11* are found in disseminated human HS ^27^, the identification and kinetic quantification of such driver mutations in the plasma are a key step in proposing and evaluating new targeted therapies in canine spontaneous model for rare and aggressive human cancer, with a double benefit in veterinary and human medicine.

Another promising application of ctDNA analysis is the follow-up of MRD during chemotherapy to assess treatment efficacy. In our pilot study, we found that ctDNA level was correlated with clinical evaluation in canine multicentric lymphomas. The 3 dogs that had a complete response also demonstrated a complete molecular response. In one dog that experienced a relapse under treatment, ctDNA was detected 42 days before clinical relapse. This is concordant with the studies of MRD targeting circulating tumoral cells (CTC) by real time PCR. In these pioneer studies, it was shown that MRD level was correlated to clinical stage at diagnosis ^61,62^ and to clinical response ^62,63^. Moreover clinical relapse could be anticipated by molecular evaluation until 86 days ^63^ and MRD level had a prognostic value ^64,65^. Further studies are needed to explore which method (ctDNA vs CTC) is more sensitive to determine MRD levels and to perform treatment follow-up in lymphoma-affected dogs. ctDNA analysis can also be performed in other cancers, as demonstrated by Beck *et al*. Indeed, they found that ctDNA was still present one year after surgical excision of a mammary carcinoma, and this was attributable to a lung metastasis ^50^. Altogether, those findings suggest that ctDNA analysis in veterinary oncology would be an objective parameter of treatment efficacy, and would be useful in earlier recognition of relapse leading to earlier re-induction or molecule switch.

In conclusion, this study shows that several somatic alterations can be found in the plasma of dogs with histiocytic sarcoma, oral malignant melanoma and multicentric lymphoma, confirming the presence of ctDNA in those canine cancers, especially in hematopoietic malignancies. Among others, two potential applications of this minimally invasive assay were explored. First the HS signature of *PTPN11* mutations and their finding in plasma could be used as a ctDNA diagnostic test to confirm a clinical suspicion of HS, especially in case of internal/disseminated forms. Second, the search of lymphoma-specific rearrangement in plasma of affected dogs is a highly accurate and personalized method to assess treatment efficacy and to anticipate a relapse. ctDNA has numerous applications in human medicine and appears to be also a promising tool in veterinary medicine with interest for future comparative oncology studies.

## Author contributions statement

Conception and design: A. Prouteau and B. Hedan. Development of methodology: A. Prouteau, J.A. Denis, E. Cadieu,, A. Lespagnol and B. Hedan. Acquisition of data: A. Prouteau, J.A. Denis, P. De Fornel, E. Cadieu,, N. Botherel, R. Ulvé, M. Rault, A. Bouzidi, R. Francois, L. Dorso, P. Devauchelle, J. Abadie, A. Lespagnol and B. Hedan. Analysis and interpretation of data: A. Prouteau, J.A. Denis, T. Derrien, C. Kergal, P. De Fornel, and B. Hedan. Writing, review, and/or revision of the manuscript: A. Prouteau, J.A. Denis, P. De Fornel, T. Derrien, J. Abadie, C. André and B. Hedan.

## Acknowledgments

The authors acknowledge all the technicians, pathologists, veterinarians and nurses that contributed to the project (particularly Drs A. Garand, F. Allais and A-S Guillory, IGDR, Rennes), the Cani-DNA Biological Resource Center (Biosit, Rennes: http://dog-genetics.genouest.org) that provided the samples, Stéphane Dréano for sequencing facilities, Yann Le Cunff for its help in statistics. The authors acknowledge Laurent Griscom (IGDR, Rennes) who assisted them for proofreading of the manuscript.

## Data availability statement

data available in article supplementary material

## Figure Legends

**Supplementary Table 1**: Main characteristics of the 133 dogs included in the study and tested for *PTPN11* mutations (PTPN11-1 for E76K and PTPN11-2 for G503V). The mutations were searched in cfDNA extracted from plasma of all dogs by ddPCR, and in the tumors of dogs with histiocytic sarcoma (HS).

**Supplementary Table 2**: Sensitivity of the ctDNA test to diagnose HS in dogs. This sensititivy depends directly on the presence of the *PTPN11* mutation in the tumor. It seems that *PTPN11* mut is more frequent in disseminated and internal forms, and when there is an involvement of lungs/mediastinum (thoracic location). Thoracic masses were confirmed by histological examination and could be associated or not with other organs involvement. This mutation is also more frequent in bernese mountain dogs.

